# StructGuy: Data leakage free prediction of functional effects of genetic variants

**DOI:** 10.64898/2025.12.01.691563

**Authors:** Alexander Gress, Carène Benasolo, Johanna Becher, Dominique Mias-Lucquin, Roman Joeres, Sebastian Keller, Olga V. Kalinina

**Affiliations:** Helmholtz Institute for Pharmaceutical Research Saarland (HIPS), Helmholtz Centre for Infection Research (HZI), Saarbruecken, Germany; Center for Bioinformatics, Saarland University, Saarbruecken, Germany; Faculty of Medicine, Saarland University, Homburg, Germany

## Abstract

The extent to which variations in protein-coding genes affect protein function has drawn the biological machine learning community’s attention to computationally model variant effect prediction tools. Multiplexed assays of variant effects (MAVE) experiments serve as a rich data source, but cannot deliver enough data for training truly large neural-net models. Therefore, zero-shot methods, for example protein language models, have increasingly gained popularity. For these methods, MAVE results serve primarily for evaluation purposes, as exemplified by the ProteinGym benchmark. In this study, we argue that the rapidly increasing amounts of MAVE data can be used to train efficient supervised methods, presenting our new tool StructGuy, based on gradient boosting trees methodology. In contrast to other supervised methods in the field, StructGuy, thanks to its dedicated training dataset and data leakage-free training process, can predict variant effects for proteins not seen during training. To evaluate this generalization ability, we constructed a dedicated benchmark and compared StructGuy with zero-shot methods from the ProteinGym leaderboard achieving a competitive performance. Further, we demonstrate that thanks to its architecture and careful feature engineering, we are able to provide fully interpretable predictions and direct explanations of the influence of mutations on protein three-dimensional structure, which favourably differs StructGuy from zero-shot tools.

## Introduction

Computer-aided interpretation of genetic variants is one of the most important bioinformatics’ contributions to personalized medicine, with the first computational methods to this end published in 2003^1^. Recently, the field experienced a boom with the advent of machine- and deep-learning methods, with the current toll of available variant effect prediction (VEP) methods nearing 200^2–4^. While over the last two decades, these methods achieved a lot, there is still a long path ahead to their true success. The holy grail for such a prediction method would be the prediction of any type of phenotype given the genetic information of an individual living being. Such end-to-end prediction spans several levels of biological complexity^5^ that all need to be correctly interpreted, which proved to be an unsolvable problem so far. Hence, the prediction process must be divided into smaller and more approachable subtasks. Such subtasks can span from (i) predicting changes in a protein’s biochemical function based on functional data on mutants to (ii) predicting pathogenicity of a variant based on large-scale population-wide observations. Both these tasks are related to establishing a genotype-to-phenotype relationship between the protein sequence and two different phenotypes: biochemical function and pathogenicity.

Many VEP tools^6–11^ have been frequently applied to solve both of these tasks, because it was observed that the predictions of the two objectives for the same proteins were highly correlated^12^. Another group of VEP tools, unsupervised models^12–18^, are designed without having a specific task in mind, and therefore can be evaluated on both tasks. For example, the EVE model^12^ has been trained to assess the probability of a protein sequence to have arisen through natural selection by considering all protein sequences from UniProt, and was demonstrated to predict well with both functional effect of mutations and their pathogenicity. In contrast, supervised models trained specifically for one of the tasks can outperform zero-shot methods on it, but fail on the other task^12,19^. Therefore the necessity for characterization and differentiation of variant effect prediction models became clear^2^ and field-wide guidelines were created^3^.

The most common type of genetic variation that affects protein sequences are non-synonymous single-nucleotide variants^20,21^ that lead to change of an amino acid in the corresponding protein and therefore affect its function. Thus, many prediction tools focus on the prediction of effects of single amino acid variations (SAVs) on the functions of individual proteins. The effect on function can be delineated into specific types, such as binding affinity towards an interacting molecule or protein structural stability. Consequently, for each function type, the amount of experimental data available for models’ training is reduced, eventually making the numbers too low to draw statistically significant conclusions. Recently, it became possible to experimentally alleviate the data scarcity problem by employing the MAVE (multiplexed assays of variant effect) technology^22–24^. A MAVE provides an experimental readout on the change of the function of almost all possible mutants in a given range of positions of a given protein. Its data have been mostly used to fine-tune and evaluate unsupervised models^12,18,25–27^, however, supervised methods were also developed, for example Envision^19^.

Recently, a lot of MAVE experiments and corresponding predictions by VEP tools have been consolidated in a benchmark set ProteinGym^25^, which comprises three parts: data from 217 MAVEs for evaluation of unsupervised methods, the same MAVE data split for protein design tasks with supervised methods, and a dataset with 62,727 genetic variants from 2,525 proteins annotated with a clinical outcome (created by selecting clinically relevant variants with a high-quality annotation from ClinVar^28^). In its protein design part, the protein data are split in such a way that a portion of positions from one protein are retained for training and the rest are used for testing the methods. This is a reasonable setup if one wants to evaluate the methods’ applicability for protein design – when a protein of interest is modified in order to optimize its function, probably in a feedback loop between computational predictions and experimental evaluation. However, a benchmark that would allow to evaluate the capability of methods to generalize predictions between different proteins is missing from ProteinGym – a drawback that we address here.

Evaluating generalizability of VEP tools across different proteins is challenging for several reasons. Perhaps the most important one is data leakage when using a naive random training/test set split. Data leakage usually refers to illicit spill-over of information between the training and test sets that leads to inflated performance^29^. In the case of MAVEs, data leakage can occur at different levels, depending on how the split is done. The easiest type of data leakage to detect is shared mutations between training and test sets. A more subtle type of data leakage is when mutations to different amino acids at the same position are placed in the training and test sets: some positions may be so susceptible to mutations that absolutely no mutations are allowed, independent of the properties of the mutant amino acid, and thus the sheer presence of this position in the training and test sets allows the model to memorize it. Further, placing mutations at different sites of the same protein to training and test sets can also cause data leakage, since some proteins may be not very robust against mutations and many mutations in them can lead to a loss of function. In this case one can train a predictor that will use the majority vote over mutations present in the training set to assign a label to any mutation in this protein from the test set, and it will be very successful – as demonstrated by Grimm et al.^30^, who called this type of data leakage “type 2 circularity”.

There are ways to counteract data leakage: for example, for the evaluation of Envision^19^ a Leave-One-Protein-Out (LOPO) training scheme was applied, where one of nine proteins in the dataset was held out for testing, and the method was trained on the remaining eight. For larger datasets, LOPO presents two drawbacks: first, when datasets include many proteins, LOPO will lead to a very high number of splits with proportionally very small test sets; and second, when datasets include homologous proteins, the presence of them in training and test sets may lead again to data leakage due to their sequence and functional similarity. Thus, the LOPO scheme needs to be extended to account for homologs to faithfully measure generalizability. Such splitting schemes, which minimize the similarity between all samples in training and all samples in test datasets, have been suggested^31^, including our own tool DataSAIL^32^.

As any machine-learning method, VEP methods featurize their input and can use different properties of a mutation for prediction. Earlier, when the determination of protein three-dimensional (3D) structures were limited, VEP tools mainly relied on sequence-derived features. This changed as the 3D structures and models became more accessible, and particularly with the introduction of AlphaFold^33^ driving the integration of structure-derived features in the recent VEP tools^11,19,26,34–38^. However, none of these methods so far include features that are based on intermolecular interactions of the corresponding protein, since predicting complexes with other proteins, nucleic acids or small molecules still present a significant challenge even for AlphaFold-like methods^39^. Unfortunately, by ignoring such features, one ignores valuable information that can inform a VEP tool. This can be alleviated by leveraging similarity of the protein(s) of interest to protein with experimentally determined structures stored in the PDB^40^. Previously, we have developed StructMAn^41,42^, an efficient tool for computing features that can be further used in VEP models. StructMAn extracts features of two types: those related to the properties of the substituted amino acids, and those related to the structural properties of the site of the substituted amino acid. The latter category puts emphasis on potential interactions by scanning all experimentally resolved 3D structures of the protein’s homologs and their complexes and integrating them into a structural descriptor of interactions of the mutated site - a feature that makes our method unique among comparable tools.

In this study, we present StructGuy (Structure-Guided VEP tool), a novel machine-learning model based on XGBoost for predicting the impact of genetic variants on protein function. We demonstrated that it achieves a competitive performance on an independent test set by combining three unique key aspects. First, we constructed a dedicated training dataset from all available MAVE experiments suitable for training of models that can generalize to unseen proteins in a supervised machine learning setup. Second, we extensively used protein structure-based features provided by StructMAn without creating surrogate sequence-based prediction models. Third, we devised an intricate feature selection and hyperparameter optimization procedure with the goal of creating a supervised model that is able to best generalize its prediction to proteins unseen during its training.

We employed and modified the current state-of-the-art benchmark ProteinGym^25^ to demonstrate the generalizability of a supervised machine learning model on the task of predicting the effects of genetic variations on protein function. We could show that StructGuy generalizes for proteins absent in the training data as well as any unsupervised prediction model in the benchmark while maintaining the superior performance of supervised models for proteins that are present in the training data. Additionally, thanks to its architecture, StructGuy is fully explainable and can fuel mechanistic molecular hypotheses behind the observed variant effects. Thus, StructGuy combines the best of the two worlds in a VEP tool that is applicable across a wide variety of setups.

## Results

### MAVE Gold Standard training dataset

We designed the MAVE gold standard (MGS) dataset so that it can be used as a training dataset for supervised machine learning VEPs that aim to be generalizable across different proteins. The key challenge here was that different MAVE experiments yield different and often incomparable readouts. A similar problem was solved in the training of Envision^19^, where a scaling method was introduced in order to aggregate the results from multiple MAVE experiments into one dataset. Since Envision was developed, the amount of available MAVE data has vastly increased, so we implemented adjustments in the scaling and quality control steps (see Methods), but generally followed a similar procedure. We scaled all observed effects such that 1 corresponds to the wildtype and 0 corresponds to a complete loss of function. Further, a quality control on the scaled dataset based on an initial evaluation of the data preprocessing employed in Envision was performed. This quality control removes experiments that cover short proteins (less than 50 amino acids), or cover a small portion (less than 40% of the proteins’ length) of a protein, or those in which the obtained distribution of experimental values was too narrow (variance less than 0.25 after scaling). We collected all available MAVE datasets from ProteinGym^25^ (version April 2025), MaveDB^43^ (version June 2025) and Domainome^45^ (version January 2025) and applied our updated scaling and quality control, obtaining a dataset with 287 proteins and 582,307 single amino acid variants (SAVs).

### Design and training of StructGuy

Each SAV in the training set was featurized using StructMAn^41^ for the calculation of structural features and features related to properties of substituted amino acids, as well as an additional feature generation pipeline to calculate evolutionary features (various features related to conservation at the mutated position, see Methods for details).

The training process consisted of several cycles of hyperparameter optimizations that kept most hyperparameters fixed while optimizing two to eight hyperparameters using a Bayes optimization scheme (see Methods). We used DataSAIL^32^ to generate four bins of proteins minimizing the sequence identity measured by MMseqs2^46^ between different bins, while proteins with high sequence similarity would be placed into the same bin. Within each cycle of fixed hyperparameters, and taking each of the bins as a test set, we performed a four-fold cross-validation for the corresponding training set comprising the remaining three bins. Of these three bins, one is held out for early stopping and the other two are used as training data for an XGBoost model^47^. Feature selection is also done on the same bin as used for early stopping by calculating SHAP values and combining them with the experimentally measured values (see Methods for details). After feature selection, a second XGBoost model is trained on the remaining features (Figure 1B), which is applied to the left-out bin to estimate the performance for a given set of hyperparameters. This procedure is repeated four times, using a different bin as the left-out test set.

**Figure 1:**
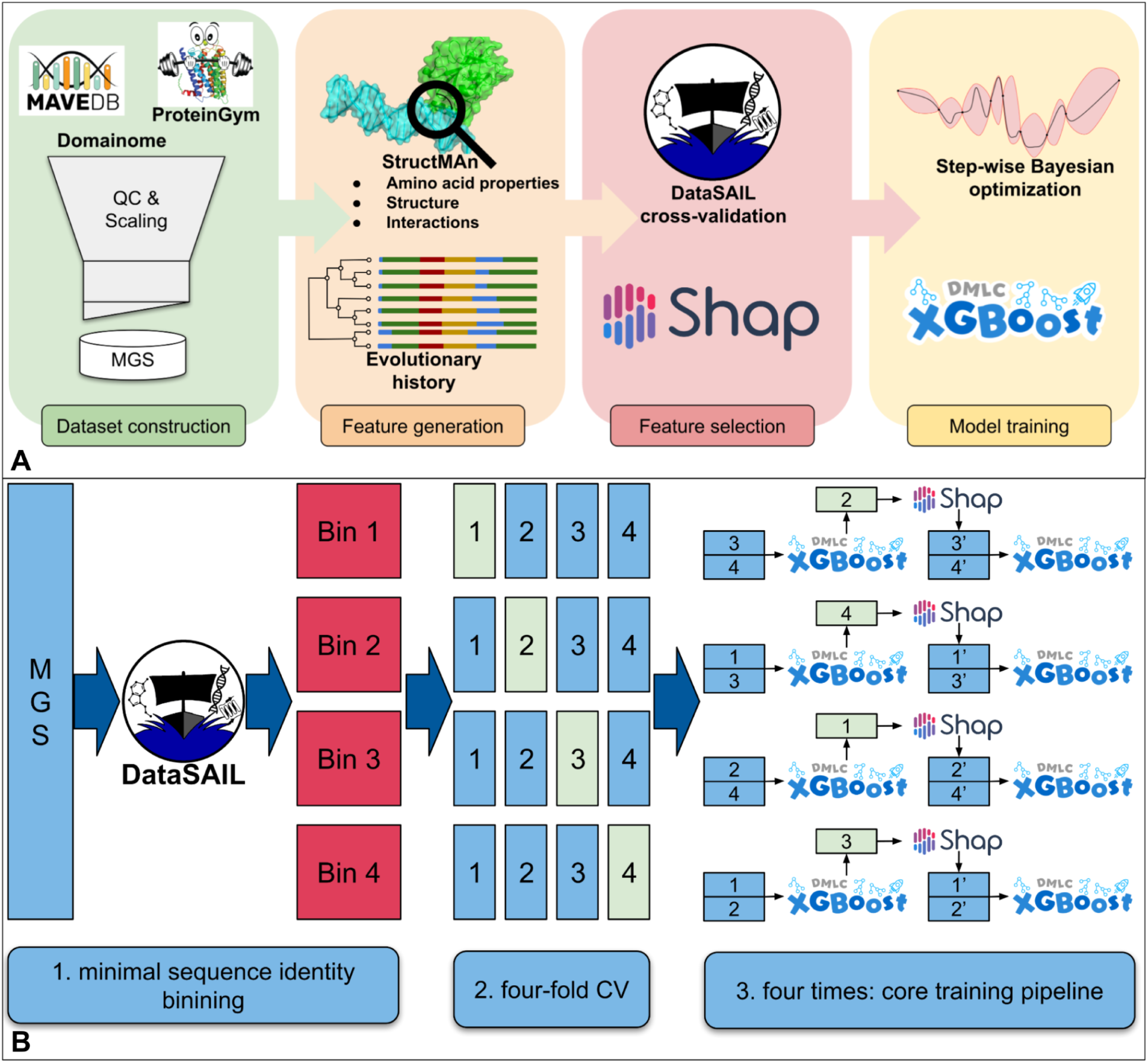
**A**. Overview of StructGuy’s training process. The MAVE gold standard (MGS) training dataset is created by filtering and rescaling MAVE data from MaveDB^43^ and ProteinGym^25^ datasets. The feature generation pipeline of StructGuy combines structural features produced by StructMAn^41^ and evolutionary features (see Methods for details). In the feature selection process, the training data are split in order to simulate prediction for unseen proteins. The model’s hyperparameters are optimized in a step-wise Bayesian optimization process. **B**. Dataset splitting scheme for the hyperparameter optimization. First, the MGS is split by DataSAIL into four bins, each of which is used for a four-fold cross validation (CV). In each CV iteration, one bin is used as held-back test data to determine the value of the objective function. Two bins are used as training data for the booster, and the remaining bin is used for early stopping and the SHAP^44^ value-based feature selection.

Once the target function does not improve anymore upon hyperparameter adjustment, the final model is trained on the full MGS dataset. It contains 7,282 boosting rounds (equal to the number of trees in the model) and a total of 240,668 nodes (parameters of the model).

### Comparison to unsupervised models from the ProteinGym benchmark

Currently, most VEP methods are unsupervised models, meaning that they were not specifically trained on the MAVE data, but simply predict some notion of sequence fitness that correlates with the MAVE outcome well, and hence they can be equally applied to any protein in the Universe. Supervised models, on the other hand, by definition have been trained on (parts of) the MAVE data, and hence if applied to the same or very similar proteins, may demonstrate an overinflated performance, which deteriorates when they are applied to proteins that are not similar to those in the training (out-of-distribution (OOD) data). This phenomenon is sometimes referred to as a poor generalizability of the model or also data leakage between the training and benchmark sets^48^. A specific evaluation is therefore necessary, testing the generalizability of supervised models, i.e. whether they can be applied to other proteins than they were trained on. The standard ProteinGym track for the evaluation of supervised models is not suitable to this end, since it was designed for evaluating models for predicting the effect of new mutations in proteins that are already present in the training set. Instead, we have modified the ProteinGym track designed for unsupervised models to enable the evaluation of supervised models for their generalizability. This is necessary in order to remove the unfair advantage that a supervised model would have due to data leakage between the training data and the benchmark set.

To this end, we constructed data leakage-free subsets of the ProteinGym dataset by placing each protein corresponding to MAVE datasets into a bin depending on its sequence identity to any of the proteins in the training set (see Methods) (Figure 2A). A typical MAVE dataset contains a wildtype protein and a set of its mutated variants, each associated with a phenotypic readout (how well it performs the biochemical function of this protein). In the MAVE datasets considered for our benchmark, we retained only sequences that differ by only one amino acid from the wildtype, omitting all multiple SAV (MSAVs) mutants from the ProteinGym dataset. While predicting MSAVs is for protein language-based prediction models basically identical to predicting SAVs, for supervised methods it would be a different task that would require the design of specialized features. As sequence identity thresholds we considered 35%, 50%, 95%, and 100% (in the latter case the benchmark contains proteins from the training set).

**Figure 2:**
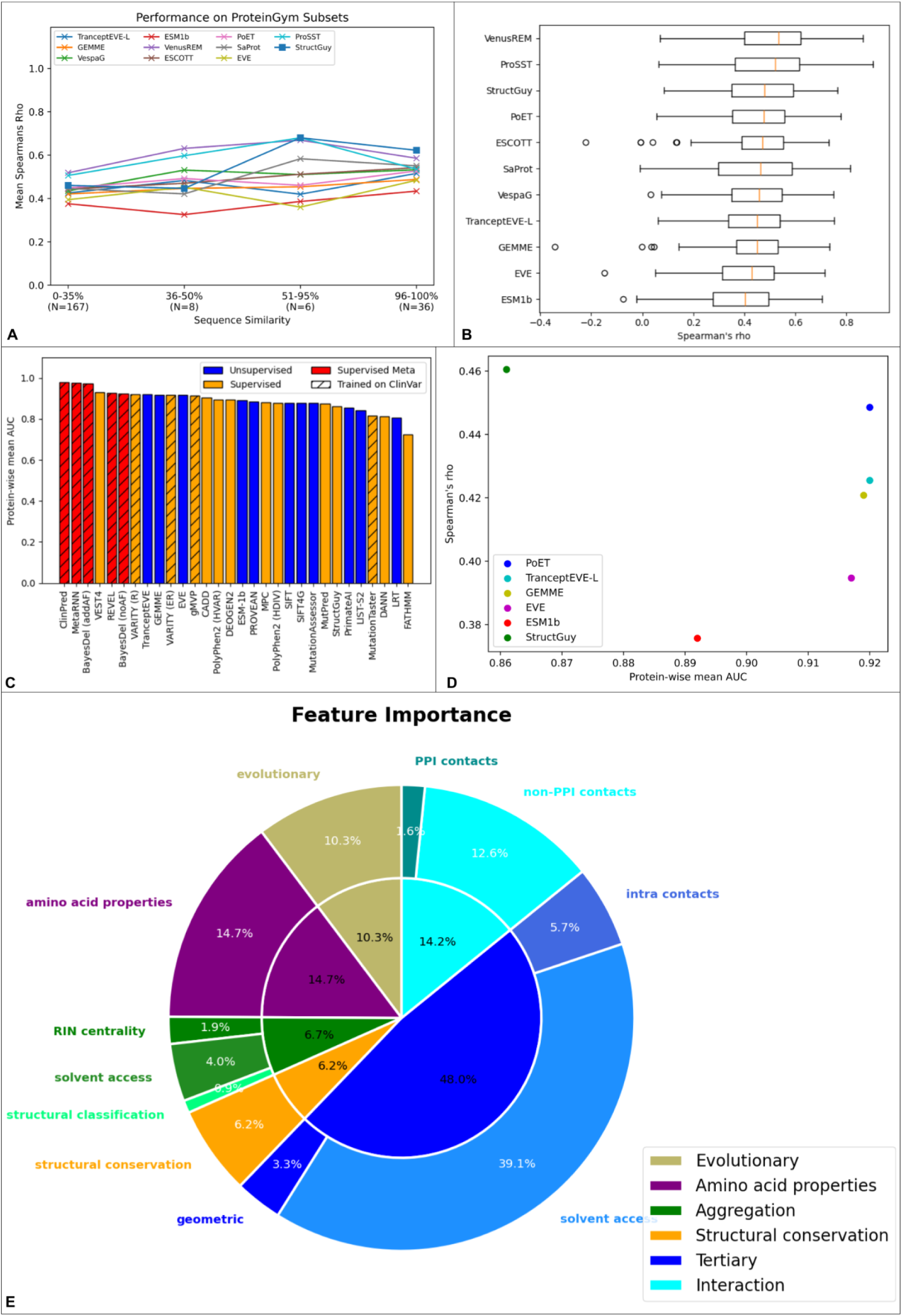
**A.** Predictive performance comparison against state-of-the-art models^9,10,12–18,49^ on the four ProteinGym-derived subsets. Similar to the metrics used in ProteinGym, each point represents the mean of function class means for one model on one subset. A function class mean is the mean Spearman’s correlation coefficient between the predicted values and measured values for all MAVEs that share the type of measured functional change (activity, binding, etc.). The left-most set of points represents a data leakage-free comparison, which are depicted in more detail in B. **B.** Boxplots for MAVE-wise Spearman’s correlation coefficient between the predicted effect values and the measured effect values (only MAVEs from the lower than 35% sequence identity subset). **C.** Comparison to the ProteinGym clinical substitution benchmark. Each bar represents the AUC score for the corresponding model predictions. The bars are colored to differentiate between types of methods and types of data the models were trained on. **D.** Concordance with clinical outcome data (X-axis) versus concordance with MAVE data (Y-axis) for VEP tools that are present in both corresponding ProteinGym tracks and StructGuy. **E.** Distribution of feature importance for different categories of features used by StructGuy.

The subset that contains only MAVEs for protein with a sequence similarity below 35% to any protein in the MGS is the largest (167 MAVE datasets) and contains over the half of all MAVEs in ProteinGym. Due to its sequence dissimilarity to our training dataset, we argue that it is fit for a fair, data leakage-free evaluation. Performance of StructGuy for this subset is very similar to the state-of-the-art, hence StructGuy can generalize as well as an unsupervised model (Figure 2B). It is important to note that at this point StructGuy cannot be fairly compared to other supervised methods using the ProteinGym benchmark, as they only have been benchmarked in the setup when the training and test sets’ mutations reside in the same proteins, hence we compare it only to unsupervised models. As sequence similarity between the benchmark proteins and the training set increases, the performance of StructGuy supersedes the state of the art. One can consider this a sign of data leakage, but this is also good news for supervised models: as more MAVE experiments that cover a larger portion of sequence space become available, their performance will increase on larger portions of the sequence space.

### Application to ProteinGym clinically relevant variants

In addition to data from MAVE experiments, ProteinGym contains a collection of clinically relevant single-amino acid substitutions and indels derived from annotated genetic variants from the ClinVar database^28^. It is constructed to benchmark the performance of supervised and unsupervised VEP models for predicting clinical outcomes. The benchmark knowingly ignores the effect of information leakage, leading to artificially inflated AUC values for methods trained on data from ClinVar (for example, ClinPred^50^). In contrast to these methods, StructGuy is trained exclusively on the MAVE data and thus is supposed to predict the effect of mutations on protein function, not the clinical outcome. Still, we applied StructGuy to the clinical outcome benchmark, assuming that clinical outcomes are strongly correlated with the functional effects of genetic variants^12^. In terms of individual variants, the overlap between the training set of StructGuy, MGS, and the ProteinGym clinical benchmark is very small (1,302 of 62,727 SAVs). Using the measured MAVE effect values as predictor for the clinical outcome for these 1302 variants, yields a modest AUC value of 0.711 demonstrating only a limited correlation between the MAVE effects and clinical outcomes and setting a baseline for assessing other methods on these data. The AUC of StructGuy on this subset is 0.713, which is to be expected since this dataset is part of its training data. Surprisingly, the AUC of StructGuy on the whole clinical substitutions dataset (Figure 2C) is much higher (0.861), but still lower than for most other methods in this benchmark, many of which were specifically trained on clinical data. This emphasizes the difference between the effect on protein function and pathogenicity and demonstrates the necessity to train dedicated tools for each task. Interestingly, several unsupervised tools (e.g. ESM-1b) that performed poorer than StructGuy on the MAVE benchmark, outperform it on the clinical data benchmark (Figure 2D), which shows that unsupervised models (and protein language models in the first place) learn a signal that correlates with effect on the whole organismal level better than with effect on precise molecular function.

### Feature importance analysis

StructGuy differs from most modern VEP tools in two key aspects: it employs a comprehensive set of protein features, including structure-based ones, and it has a gradient boosting-based classical machine learning engine. The features can be categorized into three types: *property* (features related to physicochemical properties of the substituted amino acids), *structural* (features related to protein 3D structure and the location of the mutated amino acid in it), and *evolutionary* (features related to conservation at the site of the substituted amino acid). Property features can be generated without any structural or sequence information for the particular protein. Evolutionary features are calculated using multiple sequence alignments gained by sequence similarity searches of the target protein against the UniRef90 database^51^. Structural features are calculated by analyzing protein 3D structure - experimentally resolved or predicted - of the target protein and its homologs.

We subcategorized the structural features into *tertiary, interaction, aggregation, and structural conservation.* Tertiary features are based on the tertiary structure of the target protomer without other interacting molecules, meaning that they can be computed using predicted 3D structures, for example from the AlphaFoldDB^52^. Interaction features are based on molecular interactions of the target protein that can only be observed in experimentally resolved structures retrieved from the PDB^40^. Aggregation features are calculated by analyzing a set of all 3D structures of proteins homologous to the target. They are derived from interactions of these proteins with other proteins, nucleic acids and small molecules, which may or may not be biologically relevant for the target protein. However, as it has been previously demonstrated that such interactions are well conserved among homologs^53^, we found these features to be useful for prediction (also see below). Aggregation features aggregate the structural analysis of several annotated protein structures, i.e. such features can account for cases where a protein can have different co-resolved interaction partners in different PDB entries or the accessible surface area varies between different annotated 3D structures. Structural conservation features describe the conservation of the structural microenvironment around the mutated amino acid among similar microenvironments^54^ in all PDB entries.

Feature importance analysis using the Gain metric of XGBoost for the final model trained on the full MGS training dataset reveals that structural features are the most important for prediction (Figure 2E). Structural features that cannot be calculated by only considering predicted structures (interaction, aggregation, and structural conservation) contribute more than an a quarter of the total importance of all features demonstrating the importance of experimentally resolved 3D structures and the benefit of using the features calculated from them with tools like StructMAn that integrate information from homologs. Since most such features involve describing interactions with other biological molecules in protein complexes, we argue that this type of information is indispensable for variant effect prediction.

### Case study: Peroxisome proliferator-activated receptor gamma (PPARG)

The biggest advantage of using structural features is that when combined with model interpretability techniques they can provide direct explanations for the molecular mechanism by which a mutation exerts its effect. We demonstrate this using an example of five SAVs in the human peroxisome proliferator-activated receptor gamma (PPARG), for which a MAVE dataset – measuring the activity of the protein as a transcription factor by assessing the abundance of CD36, which is controlled by PPARG – is present in ProteinGym^55^. In our benchmark dataset, this protein belongs to the part with the lowest sequence identity to the MGS, below 35%, meaning that in the training dataset of StructGuy there is no protein similar to PPARG. Yet, StructGuy was able to reproduce the measured effect reasonably well (Figure 3A) with a Spearman’s correlation coefficient of 0.615 to the target variable.

**Figure 3:**
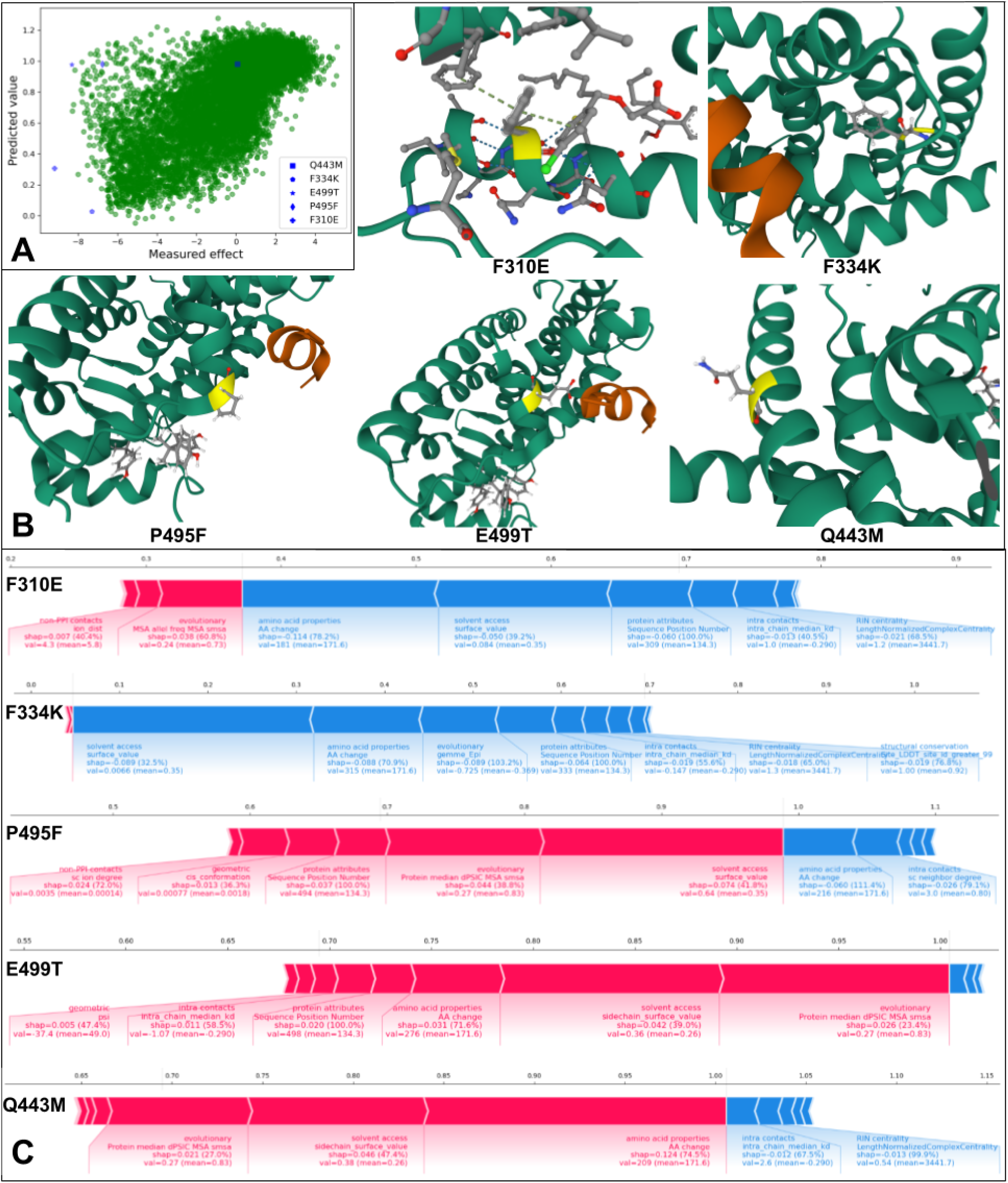
**A.** Scatterplot of predicted effect values versus measured effect values for all SAVs in the PPARG dataset^55^. The five example SAVs discussed in detail are shown in blue. **B.** The wildtype amino acids of the five example SAVs (yellow) shown inside the protein structure of PPARG (green); annotated PDB structures: F310: 6MS7, chain A, residue 282; F334: 9F7W, chain A, residue 306; P495: 9F7W: chain A, residue 467; E499: 9F7W, chain A, residue 471; Q443: 6T1S, chain A, residue 443. **C.** SHAP value force plots for the five example SAVs. The SHAP values for all features of the same feature category are summed up.

We selected three examples, where StructGuy predicted the correct outcome, two SAVs (F310E and F334K) that are disrupting the protein’s function, and one SAV (Q443M), which has a neutral effect. Additionally, we selected two example SAVs (P495F and E499T), where a strong decrease in activity was measured, yet StructGuy wrongly predicted them to have no effect. We used the StructMAn webserver^42^ to visualize the five SAVs in the protein 3D structure (Figure 3B), which can be used to give an educated guess about their mode of action. StructGuy offers another informative interpretation tool, the SHAP value force plots for individual predictions (Figure 3C).

Of all SAVs whose effect was measured in the MAVE experiment, F310E has the strongest detrimental effect. In the 3D structure, we observe a pi-pi stacking of the phenylalanine sidechain at position 310 with another phenylalanine at position 388 in PPARG and a phenol ring of the bound ligand. Removing this pi-pi stacking by introducing the negatively charged glutamic acid will destabilize the protein by destroying an intra-chain interaction of the two phenylalanines and also weakening the ligand binding. StructGuy’s decision to predict F310E as a damaging mutation is driven in the first place by statistical features (amino acid properties and protein attributes) and the solvent accessibility of the amino acid residue. Additionally, it takes into account intra-chain interactions and residue interaction network centrality of the amino acid, which together with the low solvent accessible area can be used to deduce the destabilizing effect of the mutation. The ligand interaction was not accounted for by the model’s interpretation engine.

Another functionally detrimental mutation is F334K, and here all StructGuy’s features deemed to be important point into the same direction: statistical, evolutionary and structural features all suggest that F334K will lead to a loss in protein function, but in different ways. The structural features describe Phe334 as an amino acid with high centrality that renders it important for the protein stability. Evolutionary features support this prediction by identifying the position as highly conserved and together with the statistical features render the mutation as potentially damaging.

In contrast, Q443M is a mutation in a position located on the protein surface that does not facilitate any important interaction with its side chain, at least as far as the available complexes of homologous proteins can witness. This feature is correctly identified by StructGuy as important and hence the mutation is correctly predicted as neutral.

The mechanisms of molecular effects of P495F and E499T seem to be very similar: both are a part of the interaction interface with the coactivator 1-alpha peptide (orange helix in Figure 3B), and hence both amino acids are crucial for the protein’s function, which corresponds to the MAVE measurements. However, StructGuy ignores the interaction with the peptide and focuses on the high solvent accessibility, statistical and evolutionary features of the mutations, which lead StructGuy to make wrong predictions.

## Discussion

Variant effect prediction (VEP) methods nowadays have entered a new exciting phase in their development. On the one hand, the recent and ongoing advancements in deep learning, notably large language models, led to development of new powerful unsupervised prediction models that learn biologically meaningful signals from training on all protein sequences available in databases. A typical unsupervised model (e.g. ESM-1b^13^) is trained as a masked language model and learns to predict masked amino acids in a protein sequence by extracting information from other protein sequences. In this way an unsupervised model learns which sequences are likely to occur in Nature. On the other hand, the increasing availability of large mutational (MAVE) datasets – which are still much smaller than the set of all proteins in large sequence databases, such as UniProt – laid the foundation for development of dedicated supervised models, with Envision^19^ first demonstrating the great potential of such data.

Here we present StructGuy, a novel supervised model for predicting effects of genetic variants on protein function that shows excellent capabilities to generalize to proteins unseen in the training data on the same level as state-of-the-art unsupervised prediction models. Beside a much smaller size of the training set (only 287 proteins in the MGS vs. around 250 million sequences in the UniProt, which served as the training data for e.g. ESM-1b^13^), an important advantage of training a supervised model is that it learns the target property, the change of protein function upon mutation, and not some abstract notion of sequences’ biological plausibility, as unsupervised models do. Potentially, one can combine such a supervised model with a pre-trained unsupervised model for even better results.

In this respect, it is interesting to analyze what this learned biological plausibility means in terms of effects of genetic variants on protein function and pathogenicity. In ProteinGym, five methods appear in both MAVE and pathogenicity (ClinVar) benchmarks, all of them unsupervised (Figure 2D). One cannot compare their performance on these two benchmarks in raw numbers, since one target variable is continuous (functional effect in MAVEs) and the other categorical (pathogenicity annotations in ClinVar). GEMME, which is not a machine learning model at all, but a statistical method, serves as a useful baseline in this comparison. It is evident that unsupervised models tend to improve on both tasks simultaneously. A pure masked language model ESM1b performs comparably poorly, worse than an earlier variational autoencoder EVE. Interestingly, addition of information from homologs (in PoET^14^) leads to a larger improvement on the function prediction axis. In contrast to these unsupervised models, StructGuy predicts pathogenicity not so well as they do, but excels on the functional effect axis. One can hypothesize that biological plausibility that unsupervised models learn is more related to the phenotype on the whole organismal level than in the level of individual proteins. This also agrees with the general understanding of natural selection that selects the fittest organism and not the best-performing protein in it.

Although features related to protein three-dimensional structure gain popularity both for supervised and unsupervised methods, as evidenced by an analysis of top-performing methods from ProteinGym, none of them uses structural features as extensively as StructGuy. Our feature importance analysis demonstrates that StructGuy not only takes advantage of structural features that are well known to be correlated with functional effect of mutations – for example exposure to the solvent – but also benefits from more advanced features, notably interactions in homologous proteins. Calculation of such features requires analysis of all experimentally resolved protein structures, which is computationally expensive and in our study was possible using our earlier StructMAn^41,42^ feature generation pipeline.

In contrast to most other modern VEP methods that are based on neural nets, StructGuy employs a gradient boosting technology, which is an example of tree-based ensemble methods and offers several advantages. One key advantage is the ease of prediction interpretation by identifying the contributing features. For StructGuy, this can be especially useful for explaining the molecular mechanisms, by which the genetic variants exert their effects, directly on the protein structure level, as we demonstrated in the case of PPARG. In combination with the visualization capabilities of StructMAn web server^42^, the structural features of StructGuy allow to easily construct mechanistic hypotheses of how a mutation is affecting the protein’s function. However, sometimes StructGuy misses the correct prediction by not taking into account certain important protein interactions, which requires further investigation.

Training supervised models requires special attention to the training data. It should be of a high quality and represent the intended application domain of the model well, while not overlapping with the test data to avoid data leakage. To conduct a thorough and unbiased training and evaluation of StructGuy, we created a new MAVE gold standard (MGS) dataset for training supervised models, whose utility goes beyond the training of StructGuy, which we make available to the community at https://github.com/kalininalab/StructGuy_evaluation. It can be used for the training of other supervised models in the future and will be updated as more MAVE data become available. We also constructed a new benchmark (test dataset) for the evaluation of generalization capabilities of supervised models. A model that generalizes well should be able to predict effects of genetic variants on protein function in proteins absent from training data. We assure this by minimizing data leakage between the MGS and the benchmark sets and demonstrate an excellent ability to generalize for our StructGuy model.

However, inevitably, StructGuy performs better on the proteins that are present in its training data, MGS. An ideal VEP model would perform equally well on proteins present and absent from the training data. To reach this goal, one needs to balance the model’s tendency to memorize the data with its generalization by extraction of patterns applicable to all proteins in the Universe. This can be achieved only by increasing the amount of the training data. While the number of variants in the MGS dataset is already large, the number of proteins needs to be increased.

## Methods

### Datasets

#### MAVE gold standard dataset

The MAVE gold standard (MGS) dataset is designed as a high-quality dataset to be used for training of supervised machine learning models that aim to predict effects of genetic variants on protein function. It is aimed to be as large as possible, while keeping the data quality as high as possible. The MGS is based on raw data from MAVE experiments extracted from MaveDB^43,56^, ProteinGym^25^ and Domainome^45^. Its generation includes various scaling and filtering steps, and we provide generation scripts as a fork of the MaveTools^57^ package: https://github.com/AlexanderGress/MaveTools. The full pipeline for reproduction of the dataset used in this study is provided at https://github.com/kalininalab/StructGuy_evaluation

The setup of a MAVE experiment for different proteins differs, mainly to account for the variety of protein functions. This causes read-outs of various types and on various scales. This is not an issue for unsupervised methods, since they rather rank mutants by severity of function impairment or gain; however, for supervised methods, the read-outs need to be rescaled, after which a prediction of the numerical impact on protein function is possible.

The generation pipeline starts with downloading a full copy of the MaveDB using its instance on zenodo^58^. First, all experiments that do not include protein-coding genes are filtered out. For each remaining experiment, it retrieves the amino acid sequence of the corresponding target protein. Each experiment comes with a short description and a paper abstract of the corresponding publication. We apply a keywords search in that accompanying meta data to categorize the type of the experiment into one of five classes: activity, binding, expression, stability, and organismal fitness. These five classes are the same as the experiment type annotations that are provided in ProteinGym.

In some cases experiments contain multiple score sets (sets of measurements for the same set of positions), in this case we aggregate the measurements that correspond to the same position and introduce the same variant by taking their mean. We do not aggregate different experiments that were performed on the same target protein, leading to occasional presence of identical proteins (with different Universal Resource Name (URN) identifiers) in the dataset, but with different experiment type annotation.

The list of unscaled MAVEs is supplemented by downloading all samples included in the ProteinGym substitutions track (https://marks.hms.harvard.edu/proteingym/DMS_ProteinGym_substitutions.zip). The measures from Domainome are taken from the supplementary table 5 of the corresponding publication^45^.

#### Quality control

During generation of MGS, three quality control steps are applied: a protein length filter, an SAV coverage filter, and an effect value standard deviation filter. The protein length filter excludes experiments on peptides and tiny proteins smaller than 50 amino acids. Further, some experiments are designed to measure effects in specific sites of a protein, e.g. a binding site, and thus tend to have a comparatively high amount of measurements with strong effect. Some even contain only strong effects. This leads to a skewed distribution of raw effect values and would result in spreading the strong effect scores all over the score range after scaling. To account for this, the SAV coverage filter retains only experiments that cover at least 40% of all positions of the corresponding protein. Finally, in order to learn to distinguish between individual variations, a supervised model needs to observe measurements with neutral effects as well as measurements with strong effects in the same experiment. Therefore the effect value deviation filter demands a certain level of variance in the effect values by calculating the standard deviation for all measurements of an experiment and normalizing it by the range of the interval that includes 98% of all measurements. If this normalized standard deviation is greater than 0.25, the corresponding experiment is kept in the MGS.

#### Scaling

The effect value scaling first determines three key statistics to classify the basic shape of the score distributions: a representative value of a variant with neutral (wildtype-like) effect *N*, a representative value for a strong detrimental effect *S,* and a representative value for a functionally beneficial effect *B*.

When the experiment contains measurements for synonymous mutations, the mean of their measurements is used for *N*. In other cases, the selection is based on the assumption that the majority of variants have a neutral effect^53^, and *N* is chosen to be equal to the mode of the score distribution. The mode is defined by binning the scores into 100 equally sized bins and determining the bin containing the highest number of scores. The median score of this bin is then used for *N*.

Before *S* and *B* can be calculated, one needs to determine the orientation of the scores, which is whether a detrimental effect has a higher or a lower numerical value than *N.* A proline baseline and an alanine baseline are calculated by averaging all scores for variants that replace an amino acid with a proline or an alanine, respectively, if substitutions to prolines and alanines are present in the experiment. The proline baseline represents a stronger effect compared to the alanine baseline. Otherwise, it is assumed that the average of all scores in the dataset (*A*) has a stronger effect than *N*, thus if *A>N,* then greater values mean stronger effects and vice versa. In our study, the determined orientation of effect values proved to be correct in all cases, since otherwise, we would have observed strong negative correlations for individual experiments in any cross validation setup. *S* and *B* are selected as the left and right most percentile of all scores. The scaled score is then calculated by the following formula:

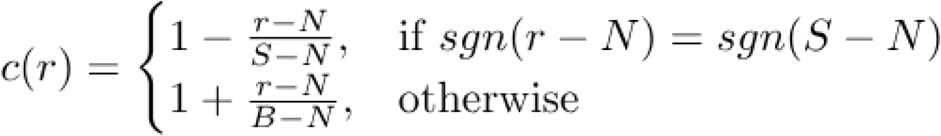

where :

*c* = scaling function

*r* = reported effect value

*S* = strong detrimental effect value

*B* = beneficial effect value

*N* = neutral effect value

#### Benchmark subsets based on sequence similarity

For the evaluation of the generalizability of StructGuy we generated dedicated benchmark datasets based on protein sequence similarity between the training and benchmark sets. The rationale for considering such datasets is twofold: 1) the amount of possible data leakage between the evaluation set and the MGS training dataset has to be minimized and 2) for all samples in the evaluation dataset, predictions from other previously published prediction models need to be present for comparison. Therefore, for the 217 MAVEs from the ProteinGym substitutions dataset, we calculated the maximal sequence identity for each of its proteins to all proteins in the MGS. Then, by applying four thresholds – 35%, 50%, 95% and 100% – we split the ProteinGym MAVEs into four subsets. A lower chosen threshold means lower data leakage between the evaluation set and training set. The subset with the lowest sequence identity threshold (35%) is the largest subset (N=167). We argue that on this subset one can provide a data leakage-free evaluation comparing StructGuy to the zero-shot prediction models listed in ProteinGym. The generation process for these evaluation subsets can be reproduced (maybe using different sequence identity thresholds) using the code at https://github.com/kalininalab/StructGuy_evaluation.

### Feature generation

The feature generation procedure of StructGuy is divided into two steps. First, all samples are processed using the structural annotation pipeline StructMAn^41,42^, which results in a table with protein structure-derived features. This table is then used as the main input for the feature generation pipeline of StructGuy, which expands it by calculating additional evolutionary features that are based on an alignment with homologous proteins.

#### Additional structural features

Most features produced by StructMAn are described in its latest publication^41^, but several new features were implemented specifically for this study. First, the relative surface area (RSA) change score is calculated, which estimates how the RSA value for the mutated amino acid differs between all available 3D structures, accessing whether solvent accessibility of the position in question may change upon changes of the conformational state of the protein.

Second, we consider a large set of features that aim to take into account the micro environment around the mutation site. To this end, MicroMiner^54^ is applied to search the PDB for structures that contain structural sites of limited size (micro environments) similar to the site that contains the variation. All annotated micro environments are grouped into bins determined by their sequence identity to the variations’ site. MicroMiner provides the root mean square deviation (RMSD) and the local distance difference test (LDDT) for each annotated microenvironment to the variations’ site. The mean RMSD and mean LDDT for each bin are used as feature values for model training.

#### Evolutionary features

Features based on the evolutionary history of the protein are frequently used for variant effect prediction. In most cases, multiple sequence alignments (MSA) are calculated for a set of proteins that share high sequence similarity to the target protein. The corresponding predicted protein structure is retrieved from the AlphaFoldDB^52^ and the MSA that was used by AlphaFold for creating that structure is fetched as well. If the MSA is not available, we apply an MMseqs2^46^ search of the protein sequence against the UniRef90 database^51^, using the resulting hits to calculate the MSA with MAFFT^59^. Then, using this MSA, features that estimate the importance of the given variation by assessing its conservation are calculated, including: GEMME scores^9^, the PSIC score^60^ of the wildtype amino acid, the PSIC score of the mutant amino acid, the dPSIC value of the variation, the median of all dPSICs of the target position, the median of dPSIC medians for a 20 position window around the target position, the median of all dPSIC medians for the whole protein, the conservation of the wildtype and the mutant amino acid, the conservation of other amino acids on the same position, and all of the previous three omitting gapped positions from the calculation.

### StructGuy training process

#### Training setup

StructGuy was trained on the MGS dataset using a four-fold cross-validation to optimize hyperparameters. First, we split the all proteins from the MGS with DataSAIL^32^ into four bins minimizing the sequence identity between proteins in different bins. In each fold, one bin is held out as a test set and training is done using the remaining three bins (Figure 1B). In the main training pipeline two of the remaining three bins are used as the inner-loop training data and one bin is used as inner-loop test data for the feature selection module of StructGuy and the early stopping routine of XGBoost^47^.

The main training pipeline consists of four steps: (1) before the training data is split, highly correlated features are removed from the dataset (see below); (2) of the three remaining bins, two are used as inner-loop training data and one as inner-loop test data. The inner-loop training data is used to fit an XGBoost model using the inner-loop test data for early stopping (the early stopping condition is that the mean protein-wise Spearman’s correlation does not change by more than 0.001 over N rounds where N is a hyperparameter); (3) This intermediate XGBoost model predicts the effect values of the inner-loop test data, and the tree SHAP values^44^ are calculated for each sample. Since, we know the true effect values for each sample in the inner-loop test set, we can calculate the loss. For each feature, the SHAP values are used to estimate the loss without that feature. Doing so, we rank features and select them by applying a threshold that is adjusted as a hyperparameter; (4) The selected features are used to train a second XGBoost model (hyperparameters are independent from the intermediate model) in the same fashion as the intermediate model. This model serves as the final model of the inner-loop.

#### Hyperparameter optimization

The hyperparameter optimization is conducted in a four-fold crossvalidation setup (Figure 1B). For each split we calculate the mean over all Spearman’s correlation coefficients for all MAVEs in the left-out test bin (*e*). We calculate the same mean for corresponding training data (*f*). Both values are used to calculate the objective function (*O*) that needs to be maximized:

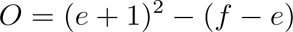

The objective function aims to optimize two properties: maximizing the test-error *e* while minimizing the difference between *f* and *e to prevent overfitting*. In total, 34 hyperparameters are optimized for StructGuy (see Supplementary Materials), which spans a 34-dimensional optimization problem, which is impossible to explore in its entirety. A common strategy to navigate this space is the Bayesian optimization procedure^61^. While it helps to find reasonable local minima, 34 dimensions span a space too vast even when individual hyperparameter ranges are limited. In the hyperparameter optimization of StructGuy we apply a step-wise Bayesian optimization approach, optimizing a small number of hyperparameters at a time, while keeping the remaining parameters fixed. StructGuy has three sets of hyperparameters: (1) two parameters that control the amount of features selected in the two feature selection steps, (2) sixteen parameters that control the XGBoost model before the second feature selection step, and (3) sixteen parameters for the main XGBoost model (see Supplementary Table 1). The target function for optimization is the mean protein-wise Spearman’s correlation.

#### Feature selection

Before the training dataset is split, highly correlated features are removed from the data. For that reason, we calculate Pearson’s correlation coefficients for all pairs of features. We iterate through all feature pairs, and if the coefficient is higher than a threshold (determined by hyperparameter optimization), one of the features is removed. Which feature is removed is determined by a score that is the product of the absolute Pearson’s correlation coefficient and the coverage of the feature (coverage is the proportion of not-none values).

The central feature selection procedure of StructGuy is designed to detect individual features that introduce data leakage into the model. Data leakage can only be estimated by testing a model on samples dissimilar to the ones used training data (out-of-distribution (OOD) samples) and therefore, the DataSAIL splits are utilized. After the training of the first XGBoost model on the training data, we use it to predict the effect scores as well as to calculate tree SHAP values (*s*) for all samples in the test data. For each feature, we calculate the feature impact (*I*) that estimates the change in loss between using the model with all features and using the model without the specific feature.

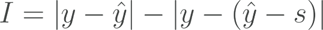

We rank all features using the mean feature impact over all samples. Then, we use a threshold, which is determined by the hyperparameter optimization, in order to select the most impactful features, which are ultimately used to train the final XGBoost model.

#### Final model training

After hyperparameter optimization, we apply the core training routine (filter highly correlated features, split training data, fit initial XGBoost model, select high impact features, fit final XGBoost model) to the full MGS training dataset. The resulting model is then called the StructGuy model.

### Evaluation

#### Comparison to unsupervised models from the ProteinGym benchmark

To evaluate StructGuy’s performance compared to unsupervised models, we downloaded all predicted values from models in ProteinGym from https://marks.hms.harvard.edu/proteingym/zero_shot_substitutions_scores.zip and calculated Spearman’s rho correlation coefficients individually for each protein, excluding MSAVs. For each group of proteins with the same measured function type, we calculate the mean over all protein-wise correlation values, and then calculate the mean over the group-wise means, which yields the final performance measure to compare different models. This procedure is identical to the one used in the ProteinGym benchmark. We repeated the same calculations for all sequence similarity threshold-based subsets (Supplementary Table 2).

#### Application to ProteinGym clinical substitutions

We downloaded the ProteinGym clinical substitutions benchmark from https://marks.hms.harvard.edu/proteingym/clinical_ProteinGym_substitutions.zip. We predicted the functional effects for all variants in this dataset using StructGuy. For each protein, we calculated the AUROC value between the predicted effect values and the clinical outcome labels from the dataset using the sklearn^62^ roc_auc_score metric.

### Reproducibility and Availability

We provide the source code of all methods required to train and apply StructGuy in a GitHub repository: https://github.com/kalininalab/structguy

Automated installation scripts that set up environments for running StructGuy are available for all linux-based systems. Additionally, a second repository contains all datasets and evaluations used in this manuscript: https://github.com/kalininalab/StructGuy_evaluation

StructGuy has been submitted to the VEP atlas: https://www.varianteffect.org/veps

## Acknowledgements

We thank Matthias Rarey and Jochen Sieg for guidance on the usage of MicroMiner. This work was partially supported by the BMFTR project Sys_CARE (project number 01ZX1908A, A.G.), the BMFTR project DrugSiderAI (project ID 031L0306A, C.B.), the HelmholtzAI project XAI-Graph (R.J.) and the Klaus Faber Foundation (O.V.K.).

